# LRP5 promotes adipose progenitor cell fitness and adipocyte insulin sensitivity

**DOI:** 10.1101/2020.03.04.976647

**Authors:** Nellie Y. Loh, Senthil K. Vasan, Manu Verma, Agata Wesolowska-Andersen, Matt J. Neville, Clive Osmond, Celia L. Gregson, Fredrik Karpe, Constantinos Christodoulides

**Author notes:** **Address for correspondence to:** Dr Constantinos Christodoulides, Oxford Centre for Diabetes, Endocrinology and Metabolism, Churchill Hospital, Oxford OX3 7LE, UK, **E-mail:**.

## Abstract

WNT signalling is a developmental pathway which plays an important role in post-natal bone accrual. We have previously shown, that in addition to exhibiting extreme high bone mass, subjects with rare gain-of-function (GoF) mutations in the WNT co-receptor LRP5 also display increased lower-body fat mass. Here, we demonstrate using human physiological studies in GoF LRP5 mutation carriers and glucose uptake assays in LRP5 knockdown (KD) adipocytes that LRP5 promotes adipocyte insulin sensitivity. We also show that a low frequency missense variant in LRP5 shown to be associated with low heel bone mineral density in a genome wide association study meta-analysis, is associated with reduced leg fat mass. Finally, using genome wide transcriptomic analyses and *in vitro* functional studies in LRP5-KD adipose progenitors (APs) we demonstrate that LRP5 plays an essential role in maintaining AP fitness i.e. functional characteristics. Pharmacological activation of LRP5 signalling in adipose tissue provides a promising strategy to prevent the redistribution of adipose tissue and metabolic sequela associated with obesity and ageing.

## INTRODUCTION

The WNT family of secreted glycoproteins play essential roles in embryonic development and adult tissue homoeostasis (1). In the canonical WNT pathway, intracellular signalling is initiated through the binding of WNT proteins to low density lipoprotein (LDL)-related protein (LRP) 5 and LRP6 co-receptors. This ultimately leads to the nuclear accumulation of the transcriptional co-activator β-catenin and WNT target gene expression (1). In addition to WNT ligands, several naturally occurring antagonists fine tune WNT signalling at the cell membrane. These include the product of the *SOST* gene sclerostin, which inhibits WNT signalling by physically interacting with LRP5 and LRP6 and preventing the binding of certain WNT ligands (2–5). WNT/β-catenin signalling is essential for normal postnatal bone accrual. Rare homozygous loss-of-function (LoF) mutations in LRP5 cause profound osteoporosis and fractures (6). This phenotype is recapitulated in mice with both germline and osteocyte-specific homozygous deletion of *Lrp5* (7). Conversely, rare heterozygous gain-of-function (GoF) LRP5 mutations that confer resistance to sclerostin binding, as well as heterozygous LoF SOST mutations, result in inherited syndromes of extreme high bone mass (HBM) (8–14).

In addition to the well-established role of LRP5 and LRP6 in skeletal homoeostasis, human and animal studies have highlighted a potential role for these receptors in systemic metabolism and adipose tissue (AT) biology. *In vitro*, knockdown (KD) of LRP5 in 3T3-L1 cells led to impaired insulin signalling and robust inhibition of adipogenesis (15). *In vivo, Lrp5*-null mice displayed impaired glucose tolerance due to reduced glucose-induced insulin secretion and developed hypercholesterolaemia following a high fat diet (HFD) (16). Furthermore, on an apolipoprotein E null background, homozygous *Lrp5* deficiency led to severe hypercholesterolaemia, impaired fat tolerance and premature and advanced atherosclerosis (17). In humans, expression of *LRP5* in AT was diminished in subjects with impaired systemic insulin sensitivity (18). Additionally, *in vitro* LRP5-KD in abdominal and gluteal adipose progenitors (APs) led to dose- and depot-dependent effects on adipogenesis (19). Furthermore, in a study of 2 pedigrees, subjects with rare LoF *LRP5* mutations had an increased prevalence of type 2 diabetes (T2D) in addition to being affected by osteoporosis (20). Finally, rare inactivating missense mutations in LRP6 were associated with autosomal dominant cardiovascular disease and features of the metabolic syndrome concomitant with osteoporosis (21 22).

We previously showed that LRP5 modulates body fat distribution. Subjects with GoF LRP5 mutations and HBM had increased lower-body fat mass whilst reciprocally, a low spine bone mineral density (BMD)-associated common intronic *LRP5* allele was associated with higher abdominal adiposity (19). Furthermore, based on *in vitro* experiments, we proposed that the effects of LRP5 on fat distribution were mediated via actions in APs. Herein, we utilised human physiological studies, *in vitro* functional assays in adipocytes and APs and genome-wide transcriptomic analyses to further dissect the role of LRP5 in AT function and glucose and lipid metabolism.

## RESEARCH DESIGN AND METHODS

### Study population

The Oxford Biobank (OBB) (23) is a population-based cohort of 30-50 year-old healthy subjects, recruited from Oxfordshire, UK. HBM *LRP5* mutation carriers were recruited from the UK-based HBM study cohort (24). Basic anthropometric data, fasting blood samples, and DXA measurements were taken (23). A sub–group underwent oral glucose tolerance tests (OGTTs). Blood was sampled at baseline (fasting) and every 30min following ingestion of 75g glucose for plasma chemistry. AT sampling and fractionation (19) are as described. All studies were approved by the Oxfordshire Clinical Research Ethics Committee. All volunteers gave written, informed consent.

### Plasma chemistry

Plasma chemistry (23) are as described. HOMA-IR, HOMA-B, and Adipo-IR were calculated as follows: HOMA-IR = fasting glucose (mmol/L) × fasting insulin (mIU/L)/22.5; and HOMA-B = [20 × fasting insulin (mIU/L)]/[glucose (mmol/L)-3.5]; Adipo-IR = fasting NEFA (mmol/L) × fasting insulin (pmol/L).

### Cell-lines

De-differentiated fat (DFAT) cells were derived by selection and de-differentiation of lipid-laden, *in vitro* differentiated immortalised human APs as described (25), with modifications.

### Doxycycline-inducible LRP5-KD cell-lines and cell culture

Oligonucleotides for shLRP5 (top: 5’CCGGGACGCAGTACAGCGATTATATCTCGAGATATAATCGCTGTACTGCGTCTTTTT; bottom: 5’AATTAAAAAGACGCAGTACAGCGATTATATCTCGAGATATAATCGCTGTACTGCGTC), and shCON (top: 5’CCGGCAACAAGATGAAGAGCACCAACTCGAGTTGGTGCTCTTCATCTTGTTGTTTTT; bottom: 5’AATTAAAAACAACAAGATGAAGAGCACCAACTCGAGTTGGTGCTCTTCATCTTGTTG) were annealed and cloned into the tet-pLKO-puro doxycycline-inducible expression lentiviral vector [kind gift of Dmitri Wiederschain (Addgene #21915)] (26). DFAT cells stably expressing tet-pLKO-puro-shLRP5 (tet-shLRP5) and tet-pLKO-puro-shCON (tet-shCON) were generated by lentiviral transduction and selection in 2μg/ml puromycin. Stable cell lines were maintained and plated under tetracycline-free conditions in DMEM-F12 supplemented with 10% FBS (ThermoFisher Scientific Gibco, #26140079), 2mM L-glutamine, 0.25ng/ml fibroblast growth factor, 100units/ml penicillin, 100μg/ml streptomycin, and 2μg/ml puromycin.

For LRP5-KD in APs, cells were cultured the following day in media containing doxycycline (or vehicle) for ~48 hours, then harvested for RNA or protein, or differentiated in the presence of doxycycline (or vehicle) throughout. Adipogenic differentiation was as previously described (19). Intracellular lipids were quantified using AdipoRed (Lonza) and a PHERAstar *FS* microplate reader (BMG Labtech).

For LRP5-KD in *in vitro* differentiated adipocytes, cells were differentiated for 13 days, then incubated in basal media (DMEM-F12 supplemented with 2mM L-glutamine, 100units/ml penicillin, 100μg/ml streptomycin, 17μM pantothenate, 33μM biotin, 10μg/ml human transferrin) containing 0.05μg/ml doxycycline (or vehicle) for 48 hours, prior to glucose-uptake assays, or harvesting for RNA and protein.

### Proliferation assays

AP proliferation was assessed in 96-well plates using CyQUANT^®^ Direct Cell Proliferation Assay (ThermoFisher Scientific) and a PHERAstar *FS* microplate reader (BMG Labtech) (19). Doubling time was calculated using the formula: *T_d_* = (*t_2_ – t_1_*) x [log (2) ÷ log (*q_2_* ÷ *q_1_*)], where *t* = time (days), *q* = fluorescence intensity (surrogate for cell number).

### Clonogenic potential

Single cells were FACS sorted into 96-well plates (one cell/well) and cultured for 15 days in conditioned media containing 0.1μg/ml doxycycline or vehicle. Cell density was assessed using CyQUANT^®^ Direct Cell Proliferation Assay and cell number estimated from a standard curve.

### Caspase-Glo 3/7 assay for apoptosis

Cells were cultured in growth media for 3 days in the presence of 0.1μg/ml doxycycline or vehicle, then a further 24 hours in serum-free media containing 0 or 100ng/ml recombinant human (rh) TNFα (ThermoFisher Scientific Gibco, #PHC3015), and doxycycline (0.1μg/ml) or vehicle. Apoptosis was assayed using the Caspase-Glo 3/7 assay (Promega) and a Veritas Microplate Luminometer (Turner Biosystems).

### Glucose uptake assay

*In vitro* differentiated cells were incubated in either fresh basal medium (to measure basal uptake) or basal medium containing 25nM insulin for 30 minutes at 37°C, 5%CO_2_. Glucose uptake was initiated by incubating cells for 10min at room temperature in uptake buffer [10μM 2-deoxy-D-glucose and 0.024MBq/ml 2-[^3^H]-deoxy-D-glucose in HEPES-buffered Saline]. Radioactivity was measured in liquid scintillant (Perkin Elmer) in a Beckman LS6500 Multipurpose Scintillation Counter (Beckman). Results were corrected for nonspecific diffusion (cells incubated in uptake buffer containing 10μM Cytochalasin B), and normalised to protein concentration.

### RNA-sequencing (RNA-seq)

Total RNA purification and on-column DNAseI-treatment were performed using the RNeasy Mini kit (QIAgen). RNA concentration was assessed using the NanoDrop ND-1000 (Labtech) and RNA quality using the Agilent 2100 Bioanalyzer (Agilent). RNA-seq (from 3 independent experiments), sequence annotation and normalization were performed at the Oxford Genomics Centre (Wellcome Trust Centre for Human Genetics, Oxford, UK). Differential gene expression was analysed using the edgeR R package (27). Gene-set enrichment analysis (GSEA) was performed on differentially expressed genes (DEGs) with FDR<0.05 in Metascape (28). Transcription factor binding site motif analyses of DEGs were performed using iRegulon in Cytoscape (29).

### Quantitative real-time PCR (qRT-CR) and Western blotting

qRT-PCR and Western blotting were performed using Taqman assays and standard protocols (19).

### Statistical Analysis

Statistical analyses were performed using R, Stata, SPSS and/or GraphPad. Statistical tests used are stated within figure legends and table footnotes.

### Data and Resource Availability

The data and resource generated during the current study are available from the corresponding author upon reasonable request.

## RESULTS

### Effects of LRP5 on systemic metabolism

To investigate the role of LRP5 on glucose and lipid metabolism we studied the metabolic phenotype of 6 individuals, 2 males and 4 females, from 3 pedigrees with extreme HBM secondary to rare heterozygous GoF LRP5 mutations (A242T and N198S) (**Table 1, Supplemental Table 1**). Each subject was closely age- and BMI-matched to 10 healthy volunteers. Compared to controls, HBM LRP5 mutation carriers were taller and had lower fasting glucose, reduced fasting insulin, and decreased Homeostatic Model Assessment for Insulin Resistance (HOMA-IR) and HOMA of β-cell function (HOMA-B) values. Additionally, HBM cases had lower AT insulin resistance (Adipo-IR). In a complimentary study, we also examined the metabolic profile of 23 homozygous carriers of a low-frequency missense LRP5 variant (V667M) which was robustly associated with reduced heel bone mineral density (BMD) in a genome wide association study (GWAS) meta-analysis (30) and therefore presumed to be LoF. Each subject was again almost identically matched to 10 healthy volunteers. Compared to controls, LoF LRP5 cases, had reduced height consistent with GWAS findings (31), in addition to displaying trends for higher fasting insulin, increased HOMA-IR and Adipo-IR as well as, elevated plasma triglycerides (**Table 1, Supplemental Table 2**). Finally, we performed OGTTs in the HBM cases and 8 independent controls (**Supplemental Table 3**). Whilst neither glucose nor insulin excursions during the OGTT were different between cases and controls in these studies, GoF LRP5 mutation carriers had lower non-esterified fatty acid (NEFA) levels at 30, 60 and 90 minutes post-OGTT as well as, NEFA area under the curve (**Fig.1, Supplemental Table 3**). We conclude that LRP5 positively regulates glucose and lipid metabolism. Furthermore, the increased systemic insulin sensitivity associated with GoF LRP5 mutations is driven partly via increased anti-lipolytic insulin action in AT.

**Fig.1.**
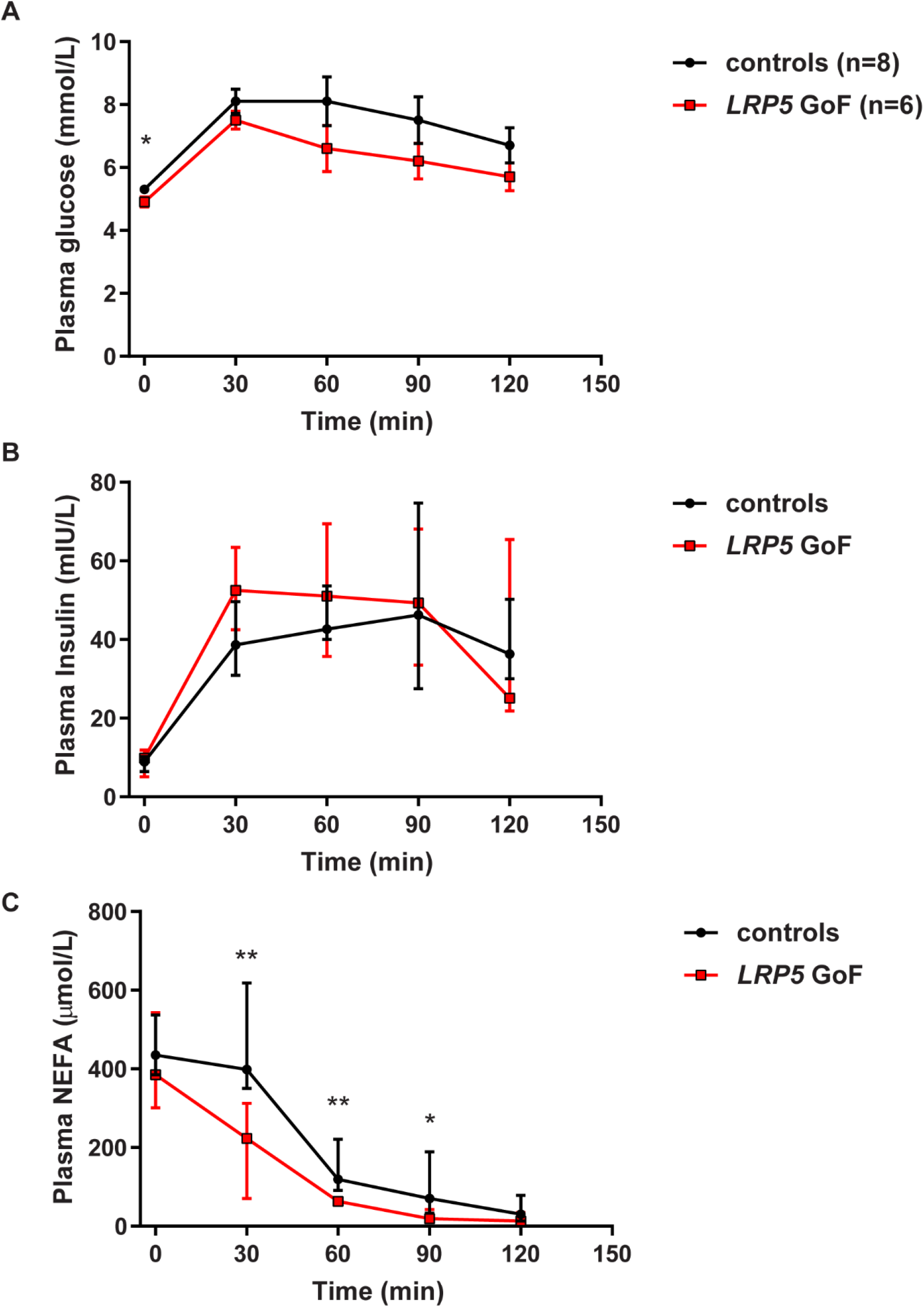
Plasma biochemical profile of control and subjects with LRP5 GoF mutation who have undergone OGTT. Plasma from blood samples collected at baseline (fasting) and at 30min intervals (up to 2h time point) following oral ingestion of glucose were measured for (**A**) glucose, (**B**) insulin, and (**C**) NEFA. Controls (n = 8), LRP5 GoF (n = 6). Graphs are mean ± SD (**A**) and median (IQR) (**B, C**). \*\**p* < 0.01, \**p* < 0.05. Statistical significance was assessed by 2-tailed unpaired Student’s t-test (**A**) and Mann-Whitney test (**B, C**).

**Table 1.** Plasma biochemical and body composition (DXA) profiles of individuals with LRP5 gain-of-function (GoF) (A242T, N198S) and LRP5 loss-of-function (LoF) (V667M) variants within cluster of age, gender and BMI matched controls

|  | **LRP5 GoF**<br>**Mean difference**<br>**(95%CI)** | **P-value** | **LRP5 LoF**<br>**Mean difference**<br>**(95%CI)** | **P-value** |
| --- | --- | --- | --- | --- |
| Gender, case (controls) | 4F, 2M (40F, 20M) |  | 15F, 8M (150F, 79M) |  |
| Age (years)* | -2.5 ± 7.8 |  | -0.7 ± 3.2 |  |
| BMI (kg/m<sup>2</sup>)* | 0.08 ± 0.42 |  | 0.01 ± 0.18 |  |
| Z-height | 0.89 (0.09, 1.69) | **0.029** | -0.45 (-0.86, -0.04) | **0.030** |
| Z-weight | 0.88 (0.08, 1.67) | **0.032** | -0.45 (-0.86, -0.04) | **0.030** |
| **DXA measurements<sup>†</sup>** |  |  |  |  |
| Z-Fat android | -0.50 (-1.31, 0.32) | 0.23 | 0.05 (-0.47, 0.57) | 0.84 |
| Z-Fat legs | 0.89 (0.09, 1.69) | **0.029** | -0.59 (-1.11, -0.06) | **0.028** |
| Z-Total fat mass | 0.45 (-0.37, 1.27) | 0.28 | -0.10 (-0.63, 0.43) | 0.71 |
| Z-Total fat percentage | -0.23 (-1.06, 0.59) | 0.57 | -0.16 (-0.69, 0.37) | 0.55 |
| Z-Android/leg fat ratio | -1.04 (-1.37, -0.71) | **2.94 x 10<sup>-7‡</sup>** | 0.43 (-0.11, 0.97) | 0.12 |
| Z-Total lean mass | 0.34 (-0.48, 1.16) | 0.41 | -0.00 (-0.53, 0.53) | 0.99 |
| Z-Lean mass legs | 0.12 (-0.71, 0.94) | 0.77 | 0.03 (-0.51, 0.56) | 0.93 |
| Z-Total BMD | 66 <sup>§</sup> | **1.25 x 10<sup>-6</sup>** | -0.70 (-1.21, -0.19) | **0.008** |
| **Biochemistry** |  |  |  |  |
| Z-glucose (fasting) | -0.97 (-1.76, -0.18) | **0.018** | -0.12 (-0.53, 0.30) | 0.58 |
| Z-insulin (fasting) | -0.92 (-1.71, -0.12) | **0.025** | 0.40 (-0.01, 0.81) | 0.06 |
| Z-HOMA-IR | -0.93 (-1.72, -0.13) | **0.023** | 0.39 (-0.02, 0.79) | 0.06 |
| Z-HOMA-B | -0.63 (-1.05, -0.22) | **0.005<sup>‡</sup>** | 0.34 (-0.07, 0.74) | 0.11 |
| Z-NEFA | -0.24 (-1.06, 0.59) | 0.56 | 0.24 (-0.17, 0.65) | 0.25 |
| Z-Adipo-IR | -0.82 (-1.24, -0.39) | **0.001<sup>‡</sup>** | 0.36 (-0.05, 0.77) | 0.09 |
| Z-triglycerides | -0.60 (-1.41, 0.22) | 0.15 | 0.38 (-0.03, 0.78) | 0.08 |

Data represents mean difference (case - control) in outcome (Z-transformed) for each case within the cluster of ten age, gender and BMI matched controls. P-value obtained from two-sample t-test (<sup>‡</sup>with Welch's correction for unequal variance) and <sup>§</sup> two-sample Wilcoxon rank-sum test.
\*represent mean and SD of the case from its controls within each cluster.
<sup>†</sup>for LoF, DXA measurements were calculated for 14 cases (10F, 4M) and their matched 139 controls on whom data was available.
Abbreviations: DXA, dual-energy X-ray absorptiometry; BMI, body mass index; CI, confidence interval; HOMA-IR, Homeostatic Model Assessment for Insulin Resistance; HOMA-B, Homeostatic Model Assessment for Insulin Secretion; NEFA, non-esterified fatty acids; Adipo-IR, adipose tissue insulin resistance.

### LRP5 cell autonomously regulates adipocyte insulin sensitivity

The *in vivo* studies outlined above highlighted that GoF LRP5 mutations were associated with enhanced AT insulin sensitivity. To determine whether LRP5 directly regulates insulin action in adipocytes, we investigated the effects of induced LRP5-KD on adipocyte insulin-stimulated glucose uptake. For these experiments we made use of a Tet-On system to express scrambled control (shCON) or LRP5 (shLRP5) shRNAs in DFAT cells generated from immortalised abdominal and gluteal adipocytes. These cells retain their depot-specific gene expression signatures and have a higher adipogenic potential than both primary and immortalised APs (32). DFAT cells were differentiated for 13 days and subsequently treated with doxycycline (dox) in hormone free media for 48 hours. Dox-treatment led to efficient (>90%) LRP5-KD in both abdominal and gluteal adipocytes (**Fig.2A-C**). LRP5-KD was not associated with changes in the expression of the master adipogenic transcription factors *PPARG* and *CEBPA* or in lipid accumulation (**Fig.2D-E**). Furthermore, no changes in the expression of mature adipocyte markers or insulin pathway genes were detected in either abdominal or gluteal LRP5-KD cells (**Supplemental Fig.1**). The only notable exception was adiponectin (*ADIPOQ*), the expression of which was decreased following LRP5-KD in both cell types (**Fig.2D**). In functional assays, LRP5-KD was associated with reduced basal glucose uptake in both abdominal and gluteal adipocytes (**Fig.2F-G**). Additionally, LRP5-KD led to impaired insulin-stimulated glucose uptake in abdominal cells (**Fig.2F**). A similar trend was detected in gluteal adipocytes (**Fig.2G**). Consistent with these findings, insulin-stimulated AKT phosphorylation was diminished in LRP5-KD abdominal DFAT adipocytes (**Fig.2H**). In complimentary experiments, *ADIPOQ* gene expression was higher in both abdominal and gluteal adipocytes from HBM cases (n = 4) *versus* controls (n = 80) (**Fig.2I**). Furthermore, a strong positive association was detected between *LRP5* and *ADIPOQ* gene expression in adipocytes from control subjects (**Supplemental Fig.2**). We conclude that LRP5 directly regulates adipocyte insulin sensitivity.

**Fig.2.**
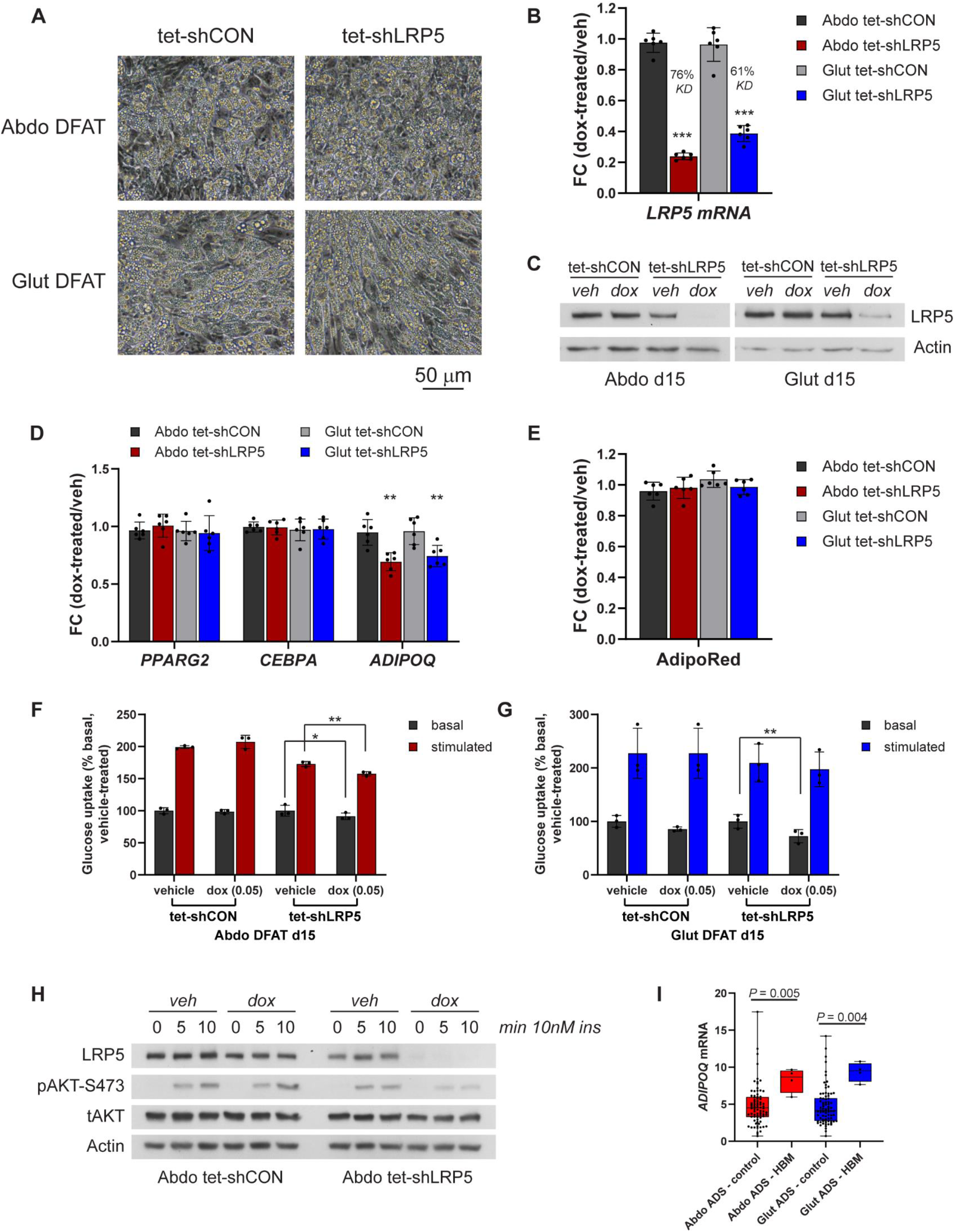
Effects of doxycycline-induced LRP5-KD on expression of adipogenic genes and on basal and stimulated glucose uptake in *in vitro* differentiated DFAT cells. (**A**) Light microscopy of abdominal and gluteal DFAT stable cell lines after 12 days adipogenic differentiation. (**B-C**) *LRP5* expression was assessed in DFAT stable cell lines at day 15 of adipogenic differentiation by (**B**) qRT-PCR (n = 6 experiments) and (**C**) Western blotting, following ~48-hour treatment with 0.05 μg/ml doxycycline or vehicle (veh) in hormone-free basal media. (**D**) qRT-PCR analyses of adipogenic genes *PPARG2*, *CEBPA* and *ADIPOQ* in *in vitro* differentiated cells from (**B**) (n = 6). (**E**) Adipogenesis, assessed by AdipoRed staining, was not different between groups (n = 6 replicates). (**F-G**) Basal and insulin-stimulated glucose uptake in *in vitro* differentiated abdominal (**F**) and gluteal (**G**) DFAT cells following ~48-hour treatment with 0.05 μg/ml doxycycline or vehicle in hormone-free basal media (n = 3 independent experiments). (**H**) Representative Western blots showing LRP5 and pAKT-S473 levels in whole cell lysates from day 15 differentiated Abdo DFAT cells following ~48-hour treatment with 0.05 μg/ml doxycycline or vehicle in hormone-free basal media, and treatment with 10 nM insulin for indicated duration. (**I**) Normalised *ADIPOQ* mRNA levels in abdominal and gluteal adipocytes (ADS) isolated from controls (43 females, 37 males) and individuals with LRP5 GoF variants (HBM; 2 females, 2 males). Analyses were adjusted for age, sex and BMI. Box and whisker plot: Whiskers are maximum and minimum values, and box represents median and interquartile range. (**B, D-G**) Histogram are means ± SD. \**p* < 0.05, \*\**p* < 0.01, \*\*\**p* < 0.001. Statistical significance was assessed by paired (**B**, **D**, **F**, **G**) and unpaired (**E**) Student’s t-test comparing doxycycline *vs*. vehicle treated groups. qRT-PCR data were normalised to *18S*. Results in **B, D, E** are expressed as fold-change (FC) of dox-treated relative to mean vehicle-treated samples.

### Effects of LRP5 on fat distribution

In addition to investigating the effects of LRP5 variants on systemic metabolism, we revisited their role on fat distribution. Compared to matched controls, GoF LRP5 mutation carriers exhibited a markedly increased total BMD but no differences in total fat mass or total lean mass (**Table 1**). Consistent with our previous report, LRP5 HBM cases also displayed increased leg and reduced android-to-leg fat mass ratio. DXA data was also available in 14 of the 23 LoF LRP5 cases. Compared to carriers of GoF LRP5 mutations homozygous V667M carriers exhibited the opposite phenotype i.e., reduced total BMD coupled with decreased leg fat mass. To gain further insights into how LRP5 modulates regional adiposity we also analysed *LRP5* expression in fractionated AT from an independent cohort of 43 females who had undergone a concomitant DXA scan to assess their fat distribution (**Table 2, Supplemental Table 4**). In age- and percent fat mass-adjusted partial correlations, both abdominal and gluteal AP *LRP5* expression correlated positively with lower-body fat mass. No correlations between mature adipocyte *LRP5* expression and regional adiposity were detected in this cohort. These data confirm and extend our previous findings, that LRP5 promotes a metabolically beneficial body fat distribution and once again highlight APs as the key AT cellular fraction mediating this effect.

**Table 2.**
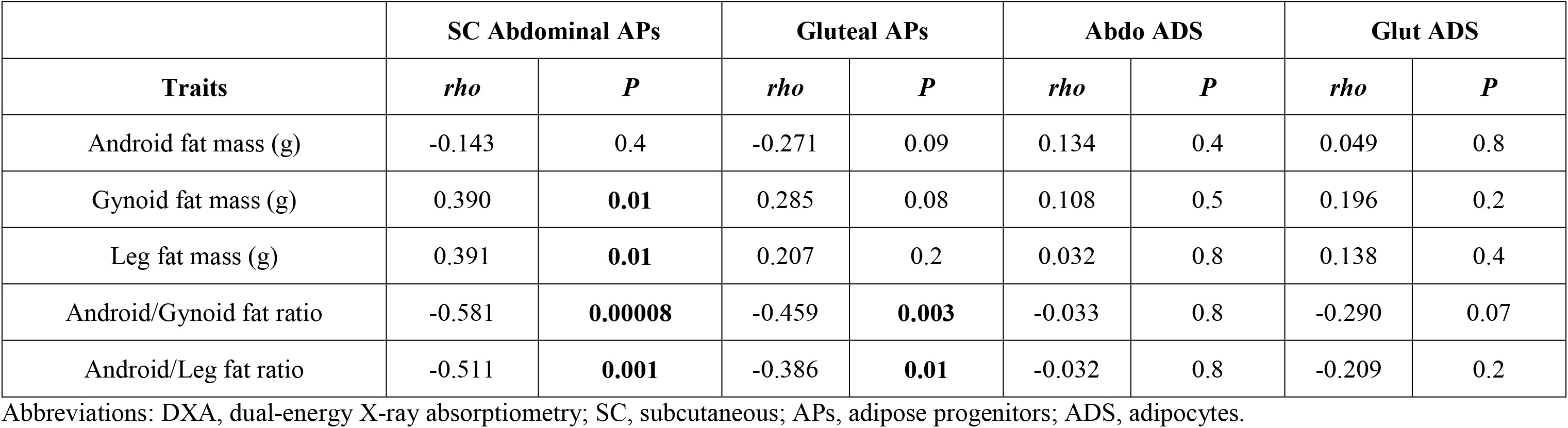
Partial correlations (Spearman’s) of measurements of body-fat distribution (DXA), with *LRP5* mRNA levels from abdominal and gluteal adipose tissue fractions from 43 women, adjusted for age, and % total fat mass.

|  | SC Abdominal APs |  | Gluteal APs |  | Abdo ADS |  | Glut ADS |  |
| --- | --- | --- | --- | --- | --- | --- | --- | --- |
| Traits | *rho* | *P* | *rho* | *P* | *rho* | *P* | *rho* | *P* |
| Android fat mass (g) | -0.143 | 0.4 | -0.271 | 0.09 | 0.134 | 0.4 | 0.049 | 0.8 |
| Gynoid fat mass (g) | 0.390 | **0.01** | 0.285 | 0.08 | 0.108 | 0.5 | 0.196 | 0.2 |
| Leg fat mass (g) | 0.391 | **0.01** | 0.207 | 0.2 | 0.032 | 0.8 | 0.138 | 0.4 |
| Android/Gynoid fat ratio | -0.581 | **0.00008** | -0.459 | **0.003** | -0.033 | 0.8 | -0.290 | 0.07 |
| Android/Leg fat ratio | -0.511 | **0.001** | -0.386 | **0.01** | -0.032 | 0.8 | -0.209 | 0.2 |

Abbreviations: DXA, dual-energy X-ray absorptiometry; SC, subcutaneous; APs, adipose progenitors; ADS, adipocytes.

### Transcriptome-wide profiling identifies biological pathways regulated by LRP5 in APs

To identify the genes and biological processes regulated by LRP5 in APs we undertook transcriptomic analyses of abdominal and gluteal DFAT cells treated with dox for 48 hours to induce LRP5-KD using RNA-seq (**Fig.3**). LRP5-KD was highly efficient in both cell types (>90%) (**Fig.3A-B**) and altered the expression of 425 genes in abdominal APs and 832 genes in gluteal cells with the majority of differentially expressed genes being down-regulated in both cell types (71% and 61% in abdominal and gluteal cells respectively) (**Fig.3C-E**). Figure 3D and 3E shows heat maps of the top 30 differentially expressed genes in abdominal and gluteal LRP5-KD cells respectively. All were down-regulated and 60% were shared between the two cell types. Gene set enrichment analysis (GSEA) revealed that the group of genes suppressed in both abdominal and gluteal LRP5-KD cells was enriched for pathways and processes involved in cell cycle, DNA repair and WNT signalling (**Fig.3F-G**). LRP5-KD in abdominal APs additionally led to down-regulation of genes involved in the apoptotic signalling pathway. Also of note, the up-regulated gene cluster in gluteal LRP5-KD cells was enriched for genes involved in cellular response to tumour necrosis factor (TNF) and apoptosis. We additionally performed transcription factor-binding site motif analysis on the promoters of genes differentially expressed in LRP5-KD APs (**Fig.3F-G**). The promoters of genes suppressed following LRP5-KD in both abdominal and gluteal cells were enriched for binding sites of multiple E2F family members as well as, FOXM1 which regulate cell proliferation. Genes whose expression was decreased in gluteal LRP5-KD cells were also enriched for TFDP1 binding sites in their promoters which co-operatively regulates cell cycle genes with members of the E2F family. Finally, genes whose expression was up-regulated in gluteal LRP5-KD cells were enriched for JUND promoter binding sites which is a component of the AP1 transcription factor complex, a key mediator of the TNF alpha-induced inflammatory response.

**Fig.3.**
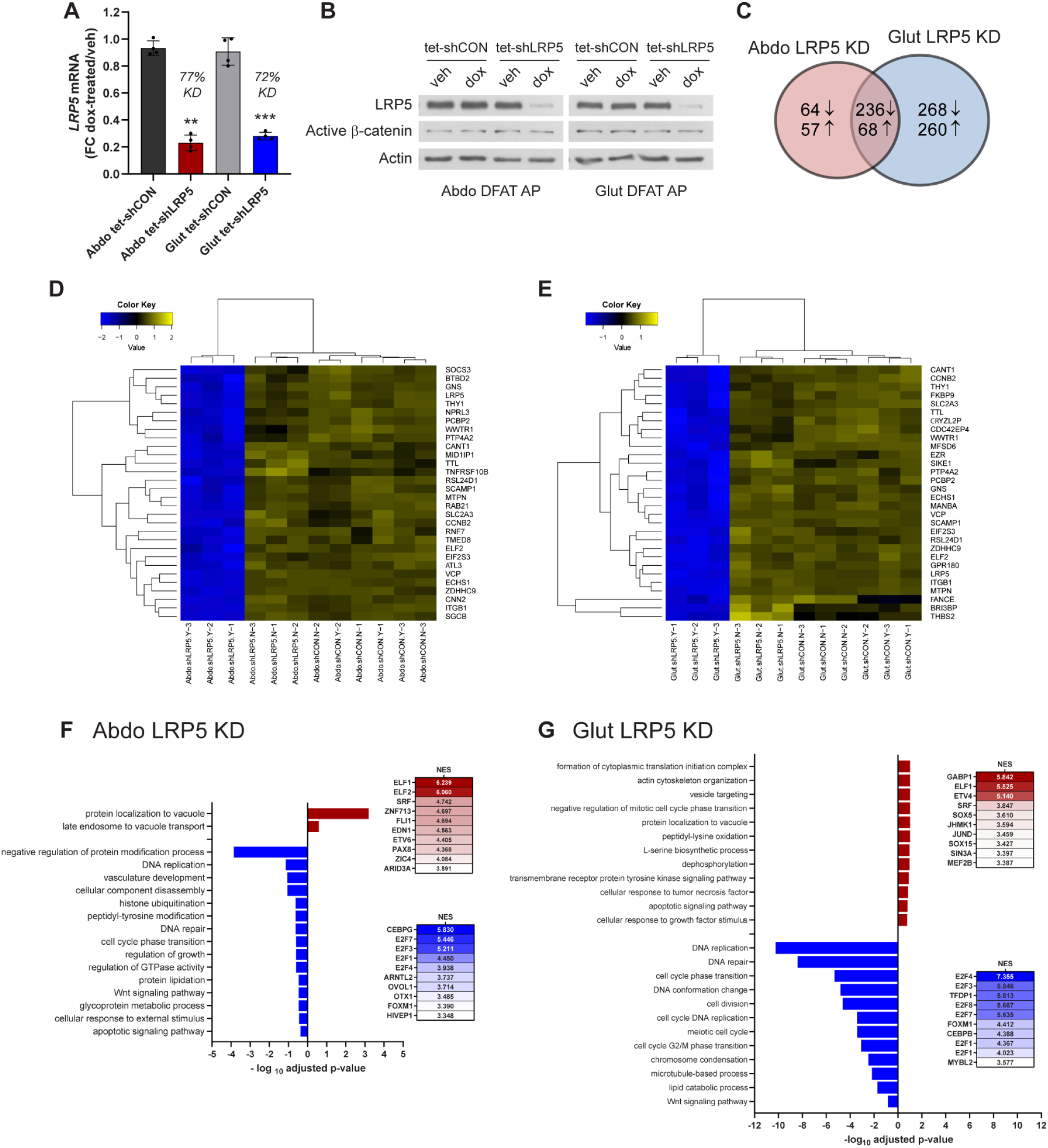
Global transcriptional profiling reveals that LRP5 regulates multiple aspects of AP biology. (**A-B**) DFAT APs, stably transduced with the tet-inducible control vector (tet-shCON) or shLRP5 vector (tet-shLRP5), were cultured in the presence of vehicle (veh) or doxycycline (final concentration of 0.1 μg/ml) for 48h to induce shRNA expression. LRP5-KD was confirmed by (**A**) qRT-PCR (n = 4 experiments) and (**B**) Western blot. qRT-PCR data were normalised to *18S* and expressed as fold-change (FC) gene expression of dox-treated samples relative to mean vehicle-treated samples. Histogram are means ± SD. (**C**) Venn diagram showing the number of significantly (FDR < 0.05) up- and down-regulated genes with doxycycline-induced LRP5-KD. (**D-E**) Heatmap showing the top 30 differentially expressed genes (DEGs) (FDR < 0.05) in doxycycline (Y) *vs*. vehicle (N) treated abdominal (**D**) and gluteal (**E**) DFAT tet-shLRP5 APs. Blue = downregulated, yellow = upregulated. (**F-G**) Pathway enrichment analyses of genes up-regulated (red) and down-regulated (blue) with doxycycline-induced LRP5-KD in (**F**) abdominal and (**G**) gluteal APs. Results of transcription factor binding-site motif analysis of DE genes, with normalized enrichment scores (NES), are shown to the right. \*\**p* < 0.01, \*\*\**p* < 0.001. Statistical significance was assessed by paired Student’s t-test comparing doxycycline *vs*. vehicle treated groups.

### LRP5-KD compromises AP fitness

To experimentally validate the results of the RNA-seq analysis we examined the functional consequences of induced LRP5-KD in DFAT APs (**Fig.4**). Consistent with the GSEA findings, LRP5-KD markedly impaired proliferation in both abdominal and gluteal APs (**Fig.4A**). No differences in the doubling time of the two cell types were detected. We next investigated the role of LRP5 in AP apoptosis (**Fig.4B**). DFAT cells were cultured in growth media with or without dox for 3 days and exposed to treatment with vehicle or TNF alpha (100 ng/ml) for the last 24 hours in serum-free media. LRP5-KD led to increased apoptosis selectively in abdominal APs independently of TNF alpha treatment. No change in Caspase Glo 3/7 activity was detected in LRP5-KD gluteal cells. Interestingly, rather than promote apoptosis, TNF alpha treatment at the dose and duration used protected abdominal APs from serum starvation-induced cell death and had no effect in gluteal cells. We additionally investigated the effects of LRP5-KD on the clonogenic potential of APs i.e. their ability to undergo “unlimited” divisions, using a novel assay which we designed in-house. Cells were FACS-sorted into single 96-well plate wells and cultured for 15 days with or without dox. LRP5-KD led to a dramatic reduction in the plating efficiency i.e. the ratio of colonies formed to number of cells seeded, in both abdominal and gluteal DFAT cells (**Fig.4C**). Furthermore, LRP5-KD cells formed much smaller colonies than their control counterparts. Finally, we revisited the effects of LRP5 on adipocyte differentiation. LRP5-KD during adipogenesis resulted in a marked and equivalent inhibition of differentiation, as determined by reduced lipid accumulation, in abdominal and gluteal cells (**Fig.4D**). We conclude that LRP5 is essential for maintaining the functional properties of APs.

**Fig.4.**
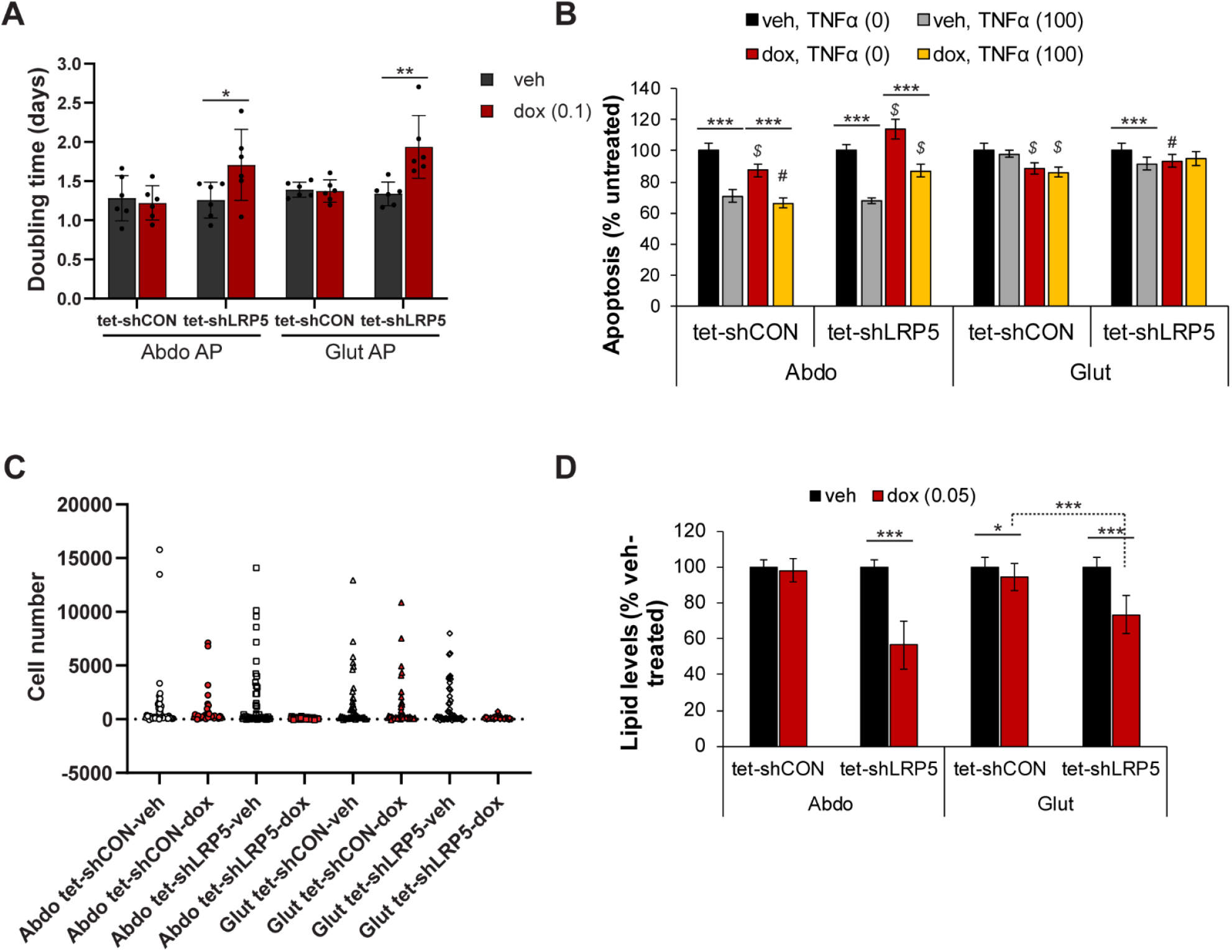
Effects of doxycycline-induced LRP5-KD on abdominal and gluteal AP biology. Effects of doxycycline-induced LRP5-KD on: (**A**) doubling time (n = 6 independent experiments; \**p* < 0.05, \*\**p* < 0.01), (**B**) apoptosis (n = 12, from three independent experiments; TNFα (0) *vs*. TNFα (100 ng/ml) treatment, \*\*\**p* ≤ 0.001; vehicle *vs*. doxycycline treatment, *^#^p* < 0.01, *^$^p* < 10^−5^), (**C**) clonogenic potential (n = 48 clones/group, representative of two independent experiments), and (**D**) differentiation (n = 18, from 3 independent experiments, \**p* < 0.05, \*\*\**p* < 0.0006). (**A, B, D**) Histogram are means ± SD. Statistical significance was assessed by 2-tailed paired (**A**) and unpaired (**B, D**) Student’s t-test.

## DISCUSSION

The present work confirms and extends our previous findings on the role of LRP5 in the regulation of systemic metabolism and fat distribution. We show for the first time that LRP5 promotes adipocyte insulin sensitivity. Additionally, based on fasting biochemistry data from 6 rare HBM cases and 60 closely matched controls (selected from a database of ~10,000 volunteers) we demonstrate that LRP5 positively regulates glucose metabolism and insulin sensitivity. Nonetheless, this latter result should be interpreted with caution as we were unable to confirm it with OGTTs albeit conducted in a small number of control subjects. Foer et al. similarly did not find any differences in glucose metabolism between 9 HBM LRP5 cases and 18 controls although, in that study LRP5 cases had lower LDL cholesterol levels (33). Unfortunately, we were unable to test this as 2 of the 6 GoF LRP5 mutation carriers were on hypolipidaemic medication. In the current study homozygous carriers of a low-frequency presumed LoF missense LRP5 variant did not display a significantly altered fasting metabolic profile, although we did detect directionally opposite trends for HOMA-IR and Adipo-IR in these subjects compared to HBM cases. This is probably because this variant, being less pathogenic than the rare GoF LRP5 mutations, exerts subtle effects on systemic metabolism. In keeping with this, Saarinen and colleagues demonstrated a high prevalence of T2D and impaired glucose tolerance in homozygous (n=2), compound heterozygous (n=1) and heterozygous (n=10) carriers of LoF LRP5 mutations associated with osteoporosis pseudoglioma syndrome (20).

Animal studies have shown that LRP5 can regulate adiposity and systemic metabolism in a cell non-autonomous manner. Mice lacking *Lrp5* in mature osteoblasts and osteocytes exhibited a progressive increase in body fat despite reduced weight coupled with reduced postnatal bone mass. Furthermore, these animals had elevated plasma triglycerides and NEFA on a chow diet and developed glucose intolerance and insulin resistance following a HFD (34 35). Opposite results were detected in chow-fed animals expressing a HBM *Lrp5* mutant allele in osteoblasts and osteocytes (34). Consistent with these findings, global *Sost* knockout mice, lacking sclerostin, displayed reduced adiposity despite normal body weight, lower plasma NEFA and increased whole-body and AT insulin sensitivity. The reverse phenotype was detected in animals with overproduction of sclerostin as a result of adeno-associated virus expression from the liver (36). In order to test whether the lower Adipo-IR in HBM subjects was due to direct or indirect effects of LRP5 in AT we generated adipocytes with inducible LRP5-KD. These experiments revealed that LRP5 can cell autonomously modulate adipocyte insulin signalling and glucose uptake. Consistent with these findings *ADIPOQ* correlated positively with *LRP5* expression in abdominal and gluteal adipocytes and was expressed more highly in adipocytes derived from HBM cases *versus* healthy controls. Furthermore, LRP5-KD in adipocytes was associated with reduced *ADIPOQ* mRNA levels. In this respect, hypoadiponectinaemia was shown to be associated with post-receptor insulin resistance (37). The mechanism whereby LRP5 promotes insulin action in adipocytes is unclear and should be the focus of future research. Work by the Kahn laboratory showed that insulin signalling is compromised in LRP5-KD 3T3-L1 cells and that the insulin receptor and LRP5 interacted in both an insulin and WNT inducible manner (15). In contrast, Kim at al. demonstrated that *Lrp5* deficient osteoblasts were insulin resistant due to the intracellular accumulation of diacylglycerol species (35). Our results do not exclude the possibility that the effects of LRP5 on whole-body metabolism could be partially mediated by actions in the skeleton e.g. via altered secretion of the insulin sensitising hormone osteocalcin (38). Given the lack of differences in total fat mass between HBM cases and controls however, it is unlikely that they involve differential energy partitioning between bone and AT. Finally, it is likely that the healthier AT expansion associated with GoF LRP5 mutations (see below) also contributes to the enhanced adipocyte insulin sensitivity.

We previously showed that GoF LRP5 mutations were associated with lower android-to-leg fat ratio (19). Herein we confirmed and extended this finding. Using a different set of controls and formal statistical methodology based on Z-scores (a measure of the standard deviations from the mean value of a reference population) we demonstrate that GoF LRP5 mutation carriers have increased leg fat mass and lower android-to-leg fat mass ratio. Gregson et al. similarly showed that 11 individuals with LRP5 HBM (including the 6 from the current study) had increased gynoid fat mass *versus* both non-LRP5 HBM cases (n = 347) as well as their unaffected relatives (n = 200) (39). Our earlier study also showed that a common intronic low spinal BMD-associated *LRP5* single nucleotide variation (SNV) (rs599083) (40) was associated with increased android-to-leg fat ratio. Based on data from GTEx however, this SNV is an expression quantitative trait locus (eQTL) for *LRP5* in lower leg skin but not leg AT. Consequently, we sought to extend this finding by examining the fat distribution of carriers of a missense LRP5 variant which was associated with reduced heel BMD in GWAS. In keeping with our earlier findings, these individuals had reduced leg fat mass. According to PolyPhen this variant is possibly damaging. Consistent with this prediction, V667M was associated with impaired activation of WNT signalling compared to wild-type LRP5 in luciferase assays in an osteosarcoma cell line (41). Elucidating the fat distribution phenotype of patients with osteoporosis pseudoglioma syndrome will ultimately provide definitive evidence of the effects of LoF LRP5 variants on regional adiposity.

In our earlier study we suggested that APs are the target cell mediating the beneficial effects of LRP5 on fat distribution. This conclusion was based on data showing that LRP5 was differentially expressed in the AP but not the adipocyte fraction of abdominal *versus* gluteal AT and that LRP5-KD in APs led to depot-dependent effects on proliferation and adipogenesis (19). Consistent with this notion we now demonstrate that *LRP5* expression in APs correlates selectively and positively with lower-body fat mass. To further elucidate the genes, pathways and biological processes regulated by LRP5 in APs we undertook genome-wide transcriptional profiling of abdominal and gluteal APs with induced LRP5-KD. These experiments highlighted a key role for LRP5 in promoting proliferation and WNT signalling in APs. Furthermore, they showed that LRP5-KD activates transcriptional programmes which lead to impaired DNA repair, and in gluteal cells result in increased inflammation. Functional studies confirmed and extended the results of the transcriptomic analysis by demonstrating that loss of LRP5 is likely to limit the expansion of subcutaneous AT by compromising the fitness of APs; namely their proliferation and survival capacity and adipogenic potential. Note that in these experiments LRP5-KD was not associated with depot-dependent effects on proliferation and differentiation consistent with the equivalent degree of KD achieved in abdominal and gluteal cells i.e. dose-dependent effects of LRP5 on AP biology as we previously proposed (19). Kato et al. similarly showed that the low bone mass phenotype of global *Lrp5* deficient mice was secondary to reduced osteoblast proliferation and function (42). Loss of *Lrp5* also led to a profound impairment of proliferation in mammary epithelial cells *in vitro* (43) and mammary stem cell maintenance *in vivo* (44). Collectively these data indicate that LRP5 performs similar functions in progenitor cells from diverse tissues.

In summary, we demonstrate that LRP5 promotes adipocyte insulin sensitivity and plays a critical role in maintaining the functional characteristics of APs. Pharmacologic activation of LRP5 in AT offers a promising approach to ameliorate the metabolic complications and fat redistribution associated with both obesity and ageing. We note in this respect that Romosozumab, a monoclonal antibody against sclerostin, was recently licenced by the European Medicines Agency for the treatment of osteoporosis in post-menopausal women and men at increased fracture risk. It will be important to determine if Romosozumab treatment has concomitant beneficial effects on metabolic health and fat distribution.

## Aknowlegements

We are grateful to the Oxford Biobank volunteers, technicians and the nurses at the Clinical Research Unit, in particular Mrs Jane Cheeseman, for their help recruiting volunteers and sample collection, as well as to Dr Toryn Poolman for his help with R. The OBB and Oxford BioResource are funded by the NIHR Oxford Biomedical Research Centre (BRC). C.C. is funded by a British Heart Foundation Clinical Research Fellowship (FS/16/45/32359). F.K. is funded by a British Heart Foundation program grant (RG/17/1/32663). We would also like to acknowledge funding support from the National Institute for Health Research (NIHR) [Oxford NIHR BRC (Diabetes & Metabolism Theme) (IS-BRC-1215-20008)], and the European Foundation for the Study of Diabetes. C.L.G. received funding for the HBM study from The Wellcome Trust (080280/Z/06/Z) and Versus Arthritis (20000). The views expressed are those of the authors and not necessarily those of the NIHR or the Department of Health and Social Care. Dr. Costas Christodoulides is the guarantor of this work and, as such, has full access to all the data in the study and takes responsibility for the integrity of the data and the accuracy of the data analysis.

## Author Contributions

Conceptualisation, C.C.; Methodology, N.Y.L., C.C.; Investigation, N.Y.L., S.K.V., A.W-A., M.V., C.C., C.L.G., M.J.N., C.O.; Writing – Original Draft, N.Y.L., C.C.; Writing – Review & Editing, All authors; Funding Acquisition, C.C., F.K., Resources, C.C., F.K., C.L.G.; Supervision, C.C.

## Competing Interests

C.C. and F.K. have received research funding from NovoNordisk and Takeda.

