## Supplementary material for "LRP5 promotes adipose progenitor cell fitness and adipocyte insulin sensitivity": Suppl. Fig.1, Suppl. Fig. 2, Suppl. Tables 1-4

**SUPPLEMENTAL FIGURES AND TABLES**

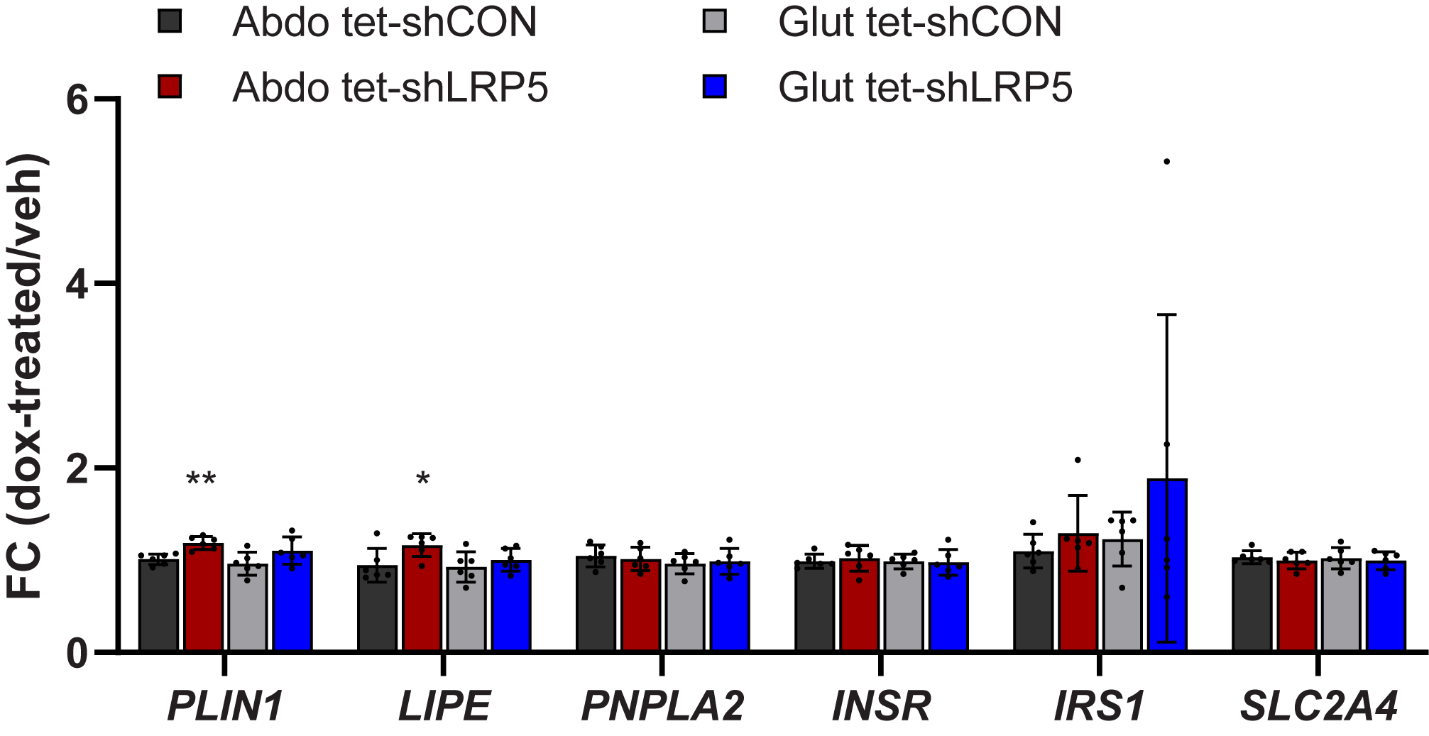

**Supplemental Fig. 1.** **mRNA expression analyses of adipogenic and insulin signalling pathway genes in *in vitro* differentiated DFAT tet-shCON and tet-shLRP5 stable cell lines.** DFAT cells were harvested for RNA at day 15 of adipogenic differentiation following ~48 hours treatment with 0.05 µg/ml doxycycline or vehicle in hormone-free basal media (n = 6 experiments). qRT-PCR data were normalised to *18S* and expressed as fold-change (FC) gene expression of dox-treated samples relative to mean vehicle-treated samples. Histogram are means ± SD. **p* < 0.05, ***p* < 0.01. Statistical significance was determined by 2-tailed paired Student’s t-test.

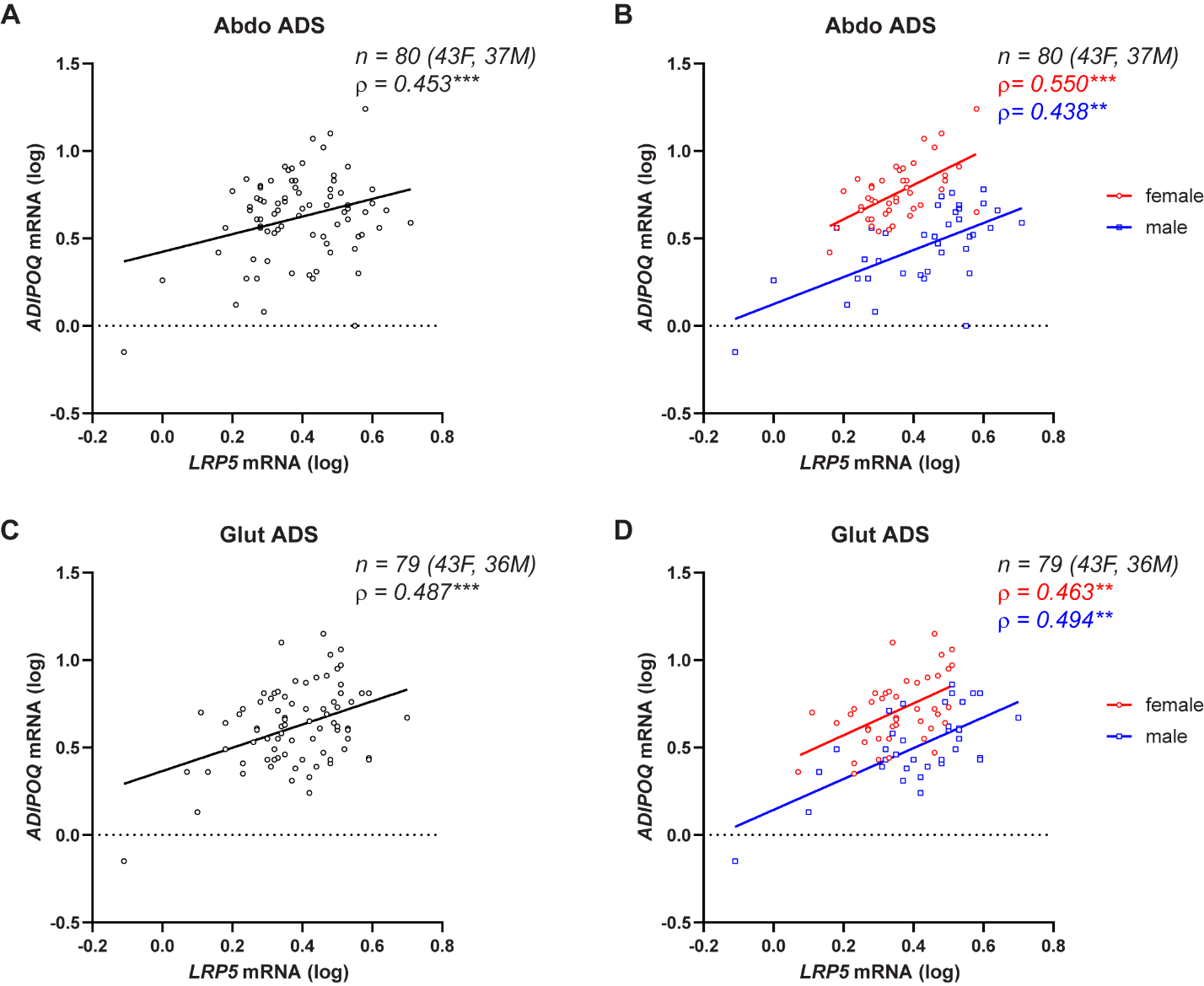

**Supplemental Fig. 2. Correlations between *LRP5* and *ADIPOQ* mRNA expression levels in human mature adipocytes**. Correlations between *LRP5* mRNA and *ADIPOQ* mRNA levels in isolated mature adipocytes (ADS) from subcutaneous abdominal (**A-B**) and gluteal (**C-D**) fat biopsies from 43 females and 37 males. (**A, C**) Non-parametric (Spearman’s) correlations for whole cohort, adjusted for age, sex and BMI. (**B, D**) Sex-specific non-parametric correlations, adjusted for age and BMI. ****p* < 0.001, ***p* < 0.01. qRT-PCR data were normalized to *18S.*

**Supplemental Table 1.** Clinical characteristics of LRP5 gain-of-function (GoF) (A242T, N198S) cases and controls

|  | **LRP5 GoF** | **Controls** | ***P*-value** |
| --- | --- | --- | --- |
|  | **(n=6)** | **(n=60)** |  |
| Sex | 4F, 2M | 40F, 20M |  |
| Age (years) | 43.0 ± 14.5 | 40.5 ± 7.4 | 0.48 |
| Height (cm) | 176.3 ± 11.2 | 170.2 ± 10.2 | 0.17 |
| Weight (kg) | 86.8 ± 19.6 | 80.4 ± 14.3 | 0.32 |
| BMI (kg/m^2^) | 27.7 ± 4.3 | 27.8 ± 4.2 | 0.97 |
| **DXA measurements** |  |  |  |
| Fat android (kg) * | 2.2 (2.0, 2.7) | 2.2 (1.8, 2.9) | 0.95 |
| Fat legs (kg) * | 10.9 (10.6, 12.4) | 9.1 (7.3, 10.8) | 0.17 |
| Total fat mass (kg) * | 30.5 (28.9, 31.8) | 27.3 (23.3, 30.2) | 0.20 |
| Total fat percentage * | 38.2 (29.3, 40.8) | 37.7 (31.1, 40.4) | 0.95 |
| Android/leg fat ratio * | 0.20 (0.16, 0.22) | 0.24 (0.20, 0.30) | **0.04** |
| Total lean mass (kg) * | 50.1 (41.2, 52.7) | 45.7 (40.4, 54.9) | 0.66 |
| Lean mass legs (kg) * | 16.3 (13.4, 20.2) | 15.7 (14.0, 18.9) | 0.86 |
| Total BMD^a^ (g/cm^2^) | 2.0 (1.8, 2.1) | 1.2 (1.1, 1.3) | **6.78 x 10^-21^** |
| **Biochemistry** |  |  |  |
| Fasting glucose (mmol/L) | 4.9 ± 0.4 | 5.3 ± 0.5 | **0.047** |
| Fasting insulin (mU/L) * | 9.9 (5.1, 11.9) | 14.4 (9.6, 18.4) | **0.041** |
| HOMA-IR * | 2.1 (1.0, 2.9) | 3.3 (2.2, 4.6) | **0.028** |
| HOMA-B * | 129 (110, 150) | 161 (127, 203) | 0.12 |
| NEFA (µmol/l) * | 385 (301, 543) | 484 (286, 617) | 0.47 |
| Adipo-IR* | 20.5 (17, 26) | 36 (21.5, 62.5) | 0.066 |
| Triglycerides (mmol/L) * | 0.7 (0.6, 1.2) | 1.0 (0.8, 1.4) | 0.16 |

Data presented as mean ± SD and *median (interquartile range) for skewed variables. *P*-values obtained from t-test and *Mann-Whitney test as applicable.

Abbreviations: BMI, body mass index; DXA, dual-energy X-ray absorptiometry; HOMA-IR, Homeostatic Model Assessment for Insulin Resistance; HOMA-B, Homeostatic Model Assessment for Insulin Secretion; NEFA, non-esterified fatty acids; Adipo-IR, adipose tissue insulin resistance.

**Supplemental Table 2.** Clinical characteristics of LRP5 loss-of-function (LoF) (V667M) cases and controls

|  | **LRP5 LoF** | **Controls** | ***P*-value** |
| --- | --- | --- | --- |
|  | **(n = 23)** | **(n = 229)** |  |
| Sex | 15F, 8M | 150F, 79M |  |
| Age (years) | 42.5 ± 6.7 | 41.8 ± 5.2 | 0.56 |
| Height (cm) | 167.2 ± 10.2 | 170.7 ± 8.8 | 0.07 |
| Weight (kg) | 72.6 ± 18.4 | 75.0 ± 15.4 | 0.49 |
| BMI (kg/m^2^) | 25.6 ± 4.1 | 25.6 ± 3.9 | 0.98 |
| **DXA measurements*** |  |  |  |
| Fat android (kg) † | 1.8 (1.2, 3.1) | 1.9 (1.2, 2.7) | 0.95 |
| Fat legs (kg) † | 6.7 (5.7, 8.6) | 8.0 (6.4, 9.7) | 0.14 |
| Total fat mass (kg) † | 21.2 (17.8, 32.7) | 23.8 (19.5, 29.3) | 0.68 |
| Total fat percentage † | 31.8 (29.6, 35.6) | 33.0 (28.3, 37.6) | 0.75 |
| Android/leg fat ratio † | 0.24 (0.15, 0.42) | 0.21 (0.16, 0.31) | 0.46 |
| Total lean mass (kg) † | 43.7 (38.3, 54.9) | 43.9 (39.6, 52.5) | 0.86 |
| Lean mass legs (kg) † | 15.5 (12.3, 20.3) | 15.2 (13.3, 17.7) | 0.95 |
| Total BMD^a^ (g/cm^2^) | 1.2 (1.1, 1.3) | 1.2 (1.1, 1.4) | **0.027** |
| **Biochemistry** |  |  |  |
| Fasting glucose (mmol/L) | 5.1 ± 0.6 | 5.1± 0.4 | 0.83 |
| Fasting insulin (mU/L) † | 13.2 (8.0, 20.1) | 11.3 (8.2, 13.8) | 0.47 |
| HOMA-IR † | 2.8 (1.7, 4.8) | 2.5 (1.9, 3.1) | 0.59 |
| HOMA-B † | 165 (116, 227) | 136 (104, 172) | 0.19 |
| NEFA (µmol/l) † | 673 (279, 813) | 494 (342, 643) | 0.19 |
| Adipo-IR † | 41 (21, 67) | 47 (31, 74) | 0.25 |
| Triglycerides (mmol/L) † | 1.0 (0.8, 1.6) | 0.9 (0.6, 1.4) | 0.17 |

Data presented as mean ± SD and †median (interquartile range) for skewed variables. *P*-values obtained from t-test and †Mann-Whitney test as applicable.

*DXA measurements available on 14 cases (10F, 4M) and their matched 139 controls.

Abbreviations: BMI, body mass index; DXA, dual-energy X-ray absorptiometry; HOMA-IR, Homeostatic Model Assessment for Insulin Resistance; HOMA-B, Homeostatic Model Assessment for Insulin Secretion; NEFA, non-esterified fatty acids; Adipo-IR, adipose tissue insulin resistance.

**Supplemental Table 3**. Clinical characteristics of subjects undergoing OGTTs

|  | **LRP5 (GoF)** | **Control** | ***P*-value** |
| --- | --- | --- | --- |
|  | **(n = 6)** | **(n = 8)** |  |
|  | 4F, 2M | 5F, 3M |  |
| Age (years) | 43.0 ± 14.5 | 45.0 ± 9.7 | 0.76 |
| Height (cm) | 176.3 ± 11.2 | 175.5 ± 7.4 | 0.87 |
| Weight (kg) | 86.7 ± 19.5 | 81.3 ± 8.1 | 0.49 |
| BMI (kg/m^2^) | 27.7 ± 4.3 | 26.4 ± 2.4 | 0.50 |
| **DXA Measurements** |  |  |  |
| Fat android (kg) | 2.2 (2.0, 2.7) | 2.4 (1.7, 2.9) | 1.00 |
| Fat legs (kg) | 10.9 (10.6, 12.4) | 9.7 (6.7, 11.9) | 0.52 |
| Total fat mass (kg) | 30.5 (28.9, 31.8) | 29.0 (23.7, 30.3) | 0.30 |
| Total fat percentage | 40.4 (31.6, 43.5) | 34.4 (29.6, 40.7) | 0.67 |
| Android/leg fat ratio | 0.20 (0.16, 0.22) | 0.26 (0.20, 0.37) | 0.12 |
| Total lean mass (kg) | 50.1 (41.2, 52.7) | 52.5 (45.4, 56.2) | 0.52 |
| Lean mass legs (kg) | 16.3 (13.4, 20.2) | 18.2 (15.3, 20.5) | 0.70 |
| Total BMD (g/cm^2^) | 2.0 (1.8, 2.1) | 1.3 (1.2, 1.3) | **8.15 x 10^-06^** |
| **Biochemsitry** |  |  |  |
| Fasting glucose (mmol/L) | 4.9 ± 0.4 | 5.3 ± 0.3 | **0.047** |
| Glucose - 30min (mmol/L) | 7.5 ± 0.7 | 8.1 ± 1.1 | 0.29 |
| Glucose - 60min (mmol/L) | 6.6 ± 1.8 | 8.1 ± 2.2 | 0.22 |
| Glucose - 90min (mmol/L) | 6.2 ± 1.4 | 7.5 ± 2.1 | 0.22 |
| Glucose - 120min (mmol/L) | 5.7 ± 1.1 | 6.7 ± 1.6 | 0.23 |
| AUC-glucose * | 775 (679, 857) | 872 (804, 959) | 0.15 |
| Fasting insulin (mU/L) * | 9.9 (5.1, 11.9) | 8.8 (6.4, 10.1) | 1.00 |
| Insulin - 30min (mU/L) * | 52.5 (42.5, 63.4) | 38.6 (30.9, 49.6) | 0.09 |
| Insulin - 60min (mU/L) * | 51.0 (35.7, 69.4) | 42.6 (40.0, 53.6) | 0.60 |
| Insulin - 90min (mU/L) * | 49.3 (33.5, 68.1) | 46.2 (27.5, 74.7) | 1.00 |
| Insulin - 120min (mU/L) * | 25.1 (21.8, 65.4) | 36.3 (30.0, 50.2) | 0.43 |
| AUC-insulin (mU/L) * | 4867 (4190, 6906) | 4668 (3935, 5279) | 0.60 |
| HOMA-IR* | 2.1 (1.0, 2.9) | 2.1 (1.6, 2.5) | 0.89 |
| HOMA-B* | 128 (113, 152) | 93 (74, 117) | 0.09 |
| Matsuda Index* | 98.1 (78.3, 149.7) | 93.1 (80.7, 125.3) | 0.79 |
| NEFA 0min (µmol/l) * | 385 (301, 543) | 435 (385, 537) | 0.51 |
| NEFA 30min (µmol/l) * | 223 (70, 312) | 398 (350, 618) | **0.009** |
| NEFA 60min (µmol/l) * | 63 (51, 65) | 119 (91, 221) | **0.008** |
| NEFA 90min (µmol/l) * | 19 (10, 42) | 70 (33, 189) | **0.038** |
| NEFA 120min (µmol/l) * | 13 (10, 34) | 30 (13, 78) | 0.27 |
| AUC-NEFA* | 16140 (12126, 20340) | 28489 (21820, 33863) | **0.005** |
| Adipo-IR * | 20 (17, 26) | 25 (19, 31) | 0.44 |
| Cholesterol (mmol/L) | 4.9 ± 0.4 | 5.0 ± 0.6 | 0.78 |
| Triglyceride (mmol/L) * | 0.72 (0.62, 1.22) | 0.71 (0.63, 1.77) | 0.81 |
| HDL-cholesterol (mmol/L) * | 1.5 (1.0, 1.7) | 1.4 (1.2, 1.5) | 0.85 |

Data presented as mean ± SD and *median (interquartile range) for skewed variables. *P*-values obtained from t-test and *Mann-Whitney test as applicable.

Abbreviations: OGTT, oral glucose tolerance test; BMI, body mass index; DXA, dual-energy X-ray absorptiometry; HOMA-IR, Homeostatic Model Assessment for Insulin Resistance; HOMA-B, Homeostatic Model Assessment for Insulin Secretion; NEFA, non-esterified fatty acids; Adipo-IR, adipose tissue insulin resistance; HDL, high density lipoprotein.

**Supplemental Table 4.** Anthropometric characteristics of female subjects from whom DXA scans and paired abdominal and gluteal AT biopsies were obtained for gene expression studies

|  | **All** | **With DXA** |
| --- | --- | --- |
| n | 61 | 43 |
| Age (years)* | 46.9 ± 1.2 (30.8, 67.4) | 50.7 ± 1.2 (37.5, 67.4) |
| BMI (kg/m^2^) * | 27.7 ± 0.5 (22.1, 40) | 27.4 ± 0.6 (22.1, 35.3) |

*mean ± SEM (min, max). Abbreviations: AT, adipose tissue; DXA, dual-energy X-ray absorptiometry; BMI, body mass index.
